# PPARδ Expression and Prognosis in Resected Pancreatic Cancer: A Retrospective Single-Centre Study

**DOI:** 10.64898/2026.09.08.750098

**Authors:** Douglas J.A. Adamson, Shaun V. Walsh, Iain S. Tait, Colin N.A. Palmer

## Abstract

**Background:** Pancreatic cancer remains one of the most lethal malignancies, with limited curative options and poor long-term outcomes. Nuclear receptors, including PPARδ, have been implicated in tumour progression.

**Methods:** Retrospectively, we analysed 25 resected pancreatic cancer specimens from patients undergoing Whipple’s procedure (pancreatoduodenectomy) between 2000 and 2009. Immunohistochemistry was performed for PPARδ. Tumour and stromal staining were assessed and correlated with clinicopathological features and survival data.

**Results:** PPARδ expression was observed in the majority of samples. This preliminary analysis suggests that PPARδ expression is associated with adverse clinicopathological features and poorer survival.

**Conclusions:** PPARδ immunohistochemistry may identify a subset of pancreatic cancers with more aggressive biology. Larger studies are warranted to validate these findings.

## Introduction

Pancreatic cancer is the 11th most common cancer worldwide, with more than 500,000 new cases annually and almost as many deaths^1^. The disease typically presents at later stages and is associated with a dismal prognosis, with fewer than 5% of patients alive at 5 years. Only a small proportion of patients are eligible for potentially curative resection^2^. Peroxisome proliferator-activated receptors (PPARs) are nuclear transcription factors involved in lipid signalling and metabolic regulation. Among them, PPARδ (also known as PPARβ/δ) has been implicated in the promotion of angiogenesis, epithelial-to-mesenchymal transition (EMT)^3^, invasion^4^, and metastasis^5^. The present study examines PPARδ expression by immunohistochemistry in a retrospective series of 25 patients who had undergone pancreatic resection and explores its relationship to outcome. By correlating PPARδ staining with clinicopathological features and survival, we aim to build on existing literature and assess its potential as a prognostic biomarker in pancreatic cancer.

## Materials and Methods

### Patients and Samples

Twenty-five patients who underwent Whipple’s resection for pancreatic cancer at our centre between 2000 and 2009. All patients were operated on with the intent of curative resection. Ethical approval for the study was obtained from the Tayside Tissue Bank.

### Immunohistochemistry

Archival formalin-fixed paraffin-embedded tissue sections were stained for PPARδ. Staining of the specimens was performed on an automated Dako immunostainer using a standard avidin-biotin-based technique. Appropriate positive and negative controls were employed. The PPARδ antibody was produced in association with our ongoing laboratory research project and was raised against the AB domain of the PPARδ protein^6^. A consultant pathologist and oncologist jointly reviewed the slides. Tumour and stromal staining were scored in both cytoplasm and nucleus on a scale of 0 (negative), 1 (weak), 2 (strong). For statistical purposes, cases were categorised as PPARδ-negative or PPARδ-positive (nuclear), or PPARδ-positive (nuclear and cytoplasmic). PPAR Tumour Cytoplasmic weak staining was assessed as “negative” as no fully negative scores were observed. The other parameters, tumour and stromal nuclear, and tumour nuclear weak and strong were collapsed to “positive”.

### Statistical Analysis

All statistical analysis was performed in STATA 13/MP. Associations between PPARδ expression and clinicopathological variables were explored using Pearson chi-square or linear regression. Survival analysis was based on follow-up which was censored at 2016 and is presented as a Kaplan Meier plot and the hazard ratios estimated using Cox regression, with and without covariates as indicated in text.

## Results

Baseline clinicopathological characteristics are summarised in Table 1. As expected for a surgical series of patients being operated on for potential curative resection, the majority of tumours were ductal adenocarcinoma of the head of pancreas, with most presenting at stage II. The median age of the patients was 67 years with a slightly higher proportion of men (60%). Tumour size measured by the assessing diagnostic pathologist ranged from 15-50mm, with the median tumour diameter being 29mm. Most tumours were graded as moderately or poorly differentiated. In this series (which predated routine pre-operative PET scanning and neoadjuvant chemotherapy), the majority had lymph node involvement on pathological assessment (88%). Just over half the series (60%) had margin positivity (“R1”) defined as being a margin of less that 1mm at any point in the specimen. The majority of patients had evidence of perineural invasion (68%) but not vascular invasion (44%).

**Table 1.** Clinicopathological characteristics of patients with resected pancreatic cancer (N = 25)

| Characteristic | n (%) or Median [Range] |
| --- | --- |
| *Demographics* |  |
| Age at resection (years) | 67 [49–78] |
| Sex (Male / Female) | 15 (60%) / 10 (40%) |
| *Tumour characteristics* |  |
| Tumour location (Head / Body / Tail) | 23 (92%) / 1 (4%) / 1 (4%) |
| Histology (Ductal adenocarcinoma / Other) | 25 (100%) / 0 (0%) |
| Tumour size (mm) | 29 [15 - 50] |
| Grade (Not specified (NS) / Well / Moderate / Poor) | 1 (4%) / 2 (8%) / 17 (68%) / 5 (20%) |
| Stage (NS / I / II / III)* | 2 (8%) / 2 (8%) / 20 (80%) / 1 (4%) |
| Lymph node involvement (Yes / No) | 22 (88%) / 3 (12%) |
| Margin status R0 [≥1mm] / R1 [$<$1mm] | 10 (40%) / 15 (60%) |
| Perineural invasion (Yes / No) | 17 (68%) / 8 (32%) |
| Vascular invasion (Yes / No) | 11 (44%) / 14 (56%) |

\* Pathological assessment using AJCC 6th edition staging.

Table 2 shows the PPARδ immunohistochemistry results and association with patient demographics and standard clinicopathological assessment factors. As most patients had tumours staged as II and in the head of the pancreas, these features were not analysed further. Age, sex, grade and LVI were not statistically associated with PPARδ positivity. Lymph node positivity, positive margins, and tumour invasion of neural tissue were all statistically associated with PPARδ expression.

**Table 2.**
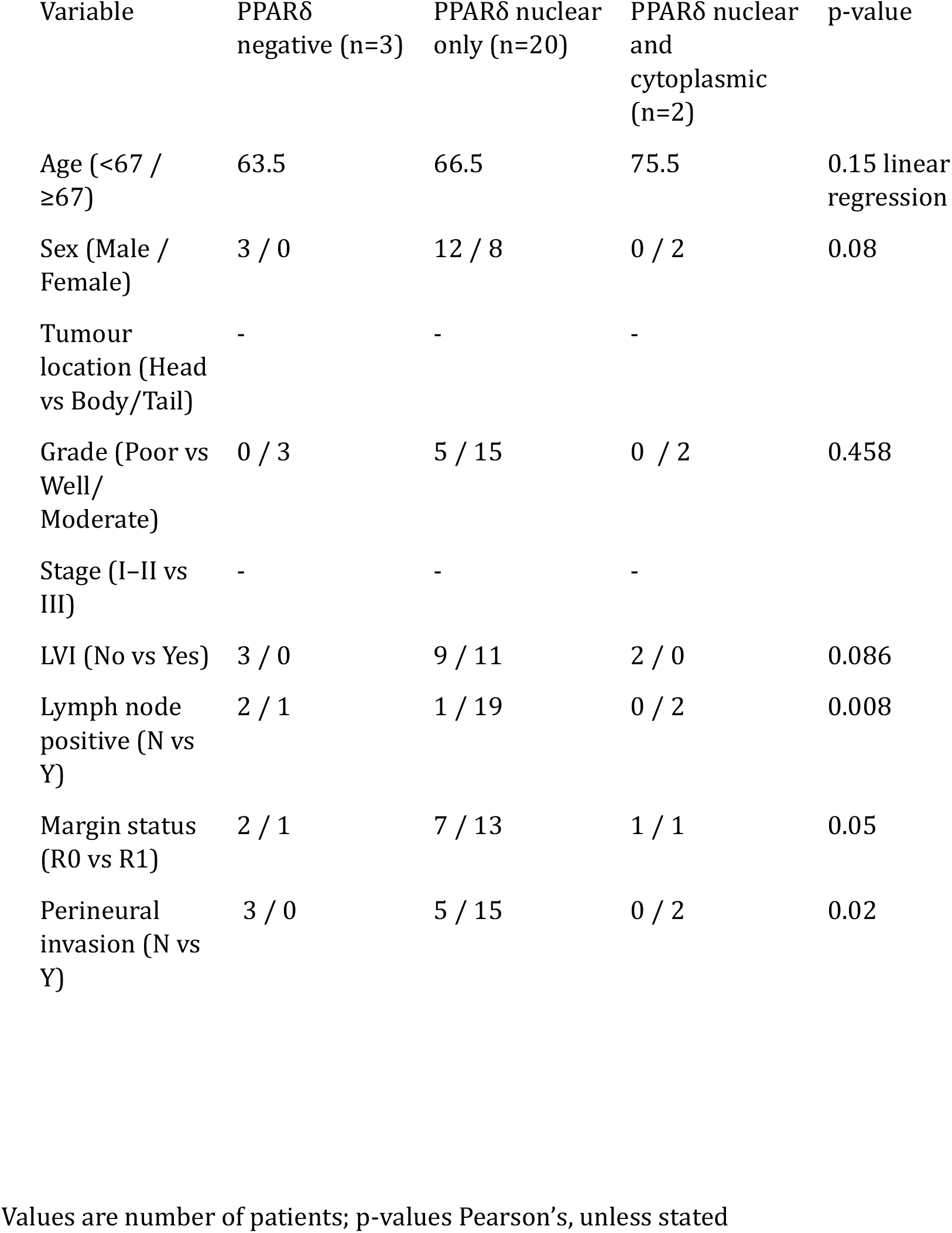
Immunohistochemistry results and association with clinicopathological variables (n = 25)

A Kaplan Meier analysis (Figure 1) was performed using time to death after surgery and a striking observation regarding the PPARδ expression was observed with expression of PPARδ in the nucleus, and expression in both nucleus and cytoplasm, demonstrating an additive risk of death with the increasingly widespread expression, with the patients having negative samples showing little progression of disease. Cox analysis revealed a Hazard ratio of 10.37 (CI 2.63-40.92), p=0.001. This may be influenced by the small negative group (n=3) and the confidence interval is wide. This was not reduced, however, when the clinical parameters that showed a relationship with PPARδ expression (node positivity, LVI, neural involvement, margin status) were included in the model, suggesting that this mortality risk is independent of these measured parameters (Adjusted HR= 18.00 (2.22-145.47), p=0.007.

**Figure 1.**
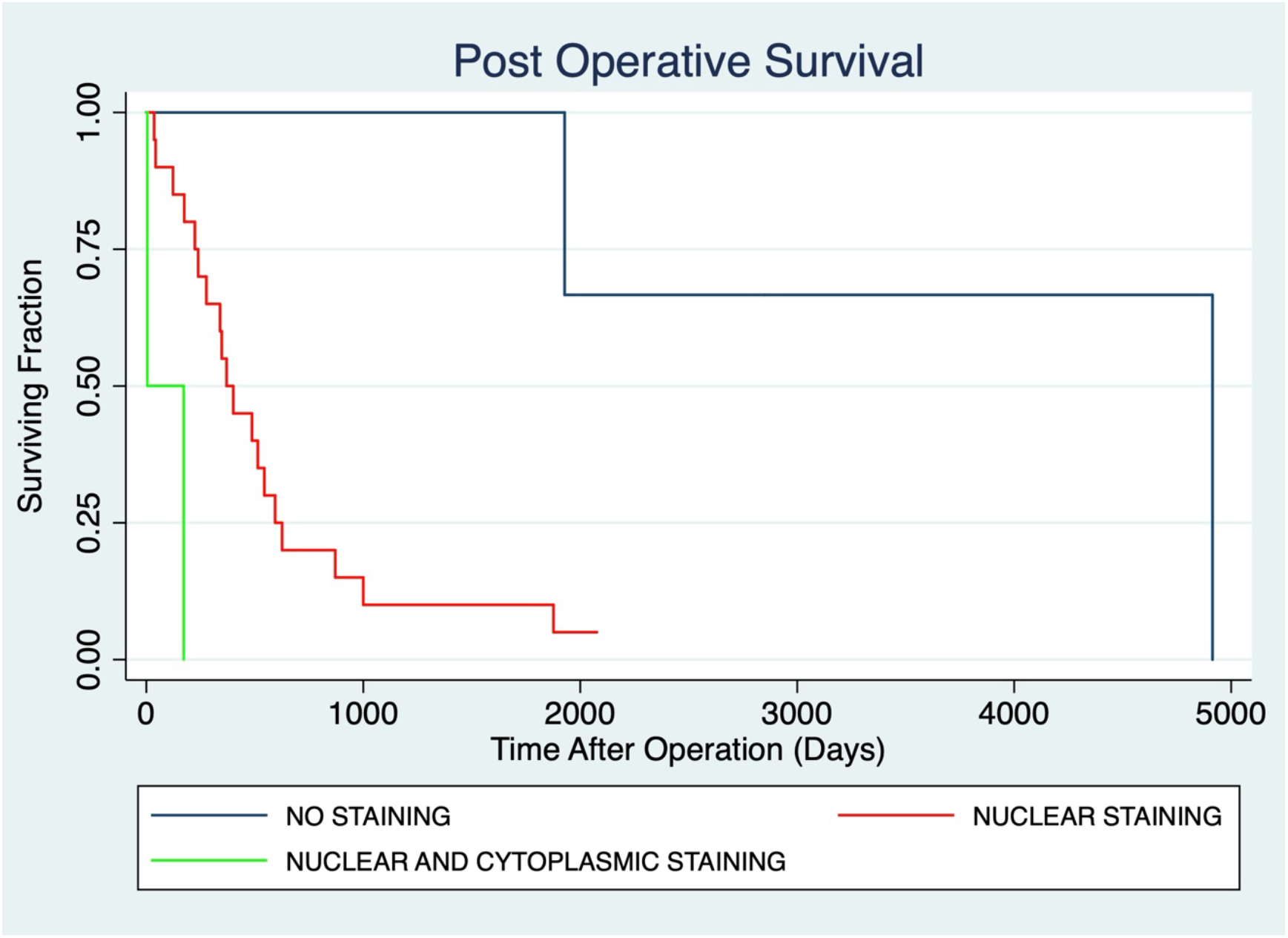
Kaplan–Meier survival curves by PPARδ expression. (All tumours with cytoplasmic staining also had nuclear staining.)

## Discussion

We chose to examine this nuclear receptor as PPARs are known to be important in lipid signalling and dietary lipid intake is significantly associated with pancreatic cancer incidence globally^7^. In addition, emerging studies suggest that nuclear receptors are important in the development and the homeostasis of the pancreas^8^. It is important to note that this is a small series of just 25 patients from a single centre undergoing pancreatoduodenectomy. The study design was retrospective, a non-standard antibody was used, and the patients were from period when pre-operative PET scanning and neoadjuvant chemotherapy were not routine, so all associations and conclusions need to be cautiously interpreted, but this analysis suggests that PPARδ expression may be an adverse prognostic marker in resected pancreatic cancer. Our preliminary findings that PPARδ expression is associated with lymph node involvement, positive margins, and perineural invasion align with prior evidence linking PPARδ with aggressive tumour biology such as invasion, angiogenesis, and immune evasion.

Experimental data suggest that PPARδ expression is upregulated in the pre-malignant pancreatic intraepithelial neoplasia (PanIN) and this is driven in part by oncogenic K-ras^9^, with potential destructive effects on the tumour immune microenvironment. The development of pancreas cancer therefore has some similarity to colon cancer development with premalignant lesions, inflammation, and different gene ‘hits’^8,10,11^. Other previous studies have highlighted the role of PPARδ in pancreatic cancer progression. For instance, PPARδ has been identified as a key hub in the angiogenic network, with its expression upregulated in pancreatic cancer tissues compared to normal pancreas. Higher PPARδ expression levels correlate with advanced pathological tumour stage, increased risk for tumour recurrence, and distant metastasis. This suggests that PPARδ contributes to the “angiogenic switch,” a critical step in tumour progression^12^. This supports our observation of PPARδ’s association with adverse clinicopathological features, suggesting it may drive vascular remodeling and metastatic potential in resected cases. More recent investigations have uncovered a novel cancer cell-intrinsic axis involving glutamic-oxaloacetic transaminase 2 (GOT2) and PPARδ, which suppresses antitumour immunity in pancreatic ductal adenocarcinoma (PDAC). It appears that GOT2 functions as a nuclear fatty acid transporter that binds to and activates PPARδ, leading to spatial restriction of CD4+ and CD8+ T cells from the tumour microenvironment, thereby promoting immune evasion and tumour growth^13^. This GOT2-PPARδ interaction extends beyond traditional metabolic roles, highlighting PPARδ’s involvement in transcriptional regulation of immunosuppressive pathways. This mechanism could explain the poorer survival trends in PPARδ-positive tumours in our cohort, as immune evasion is a key determinant of long-term outcomes in pancreatic cancer.

Our study also resonates with investigations into PPARδ’s metabolic influences. PPARδ has been linked to lipid metabolism alterations in pancreatic cancer cells. Agonist-mediated activation of PPARδ has been shown to modulate fatty acid desaturases like SCD1 and Δ6D in pancreatic cancer cells, potentially altering lipid profiles that support proliferation and invasion^14^. These findings underscore PPARδ’s potential multifaceted role in modulating metabolic and signalling pathways that support pancreatic cancer aggressiveness. PPARδ may also be associated with invasion and metastasis in pancreatic cancer—in human pancreatic tissue, levels of PPARδ and MMP-9 mRNA are raised, while BCL-6 is reduced. MMP-9 is associated with early invasive behaviour of tumour cells, with levels 10-fold higher in pancreatic cancer compared with tissue from chronic pancreatitis^15^.

While these studies used *in vitro* models, they provide a mechanistic basis for why PPARδ expression might confer a more aggressive phenotype *in vivo*, as seen in our immunohistochemical correlations. Building on these metabolic roles, Parejo-Alonso *et al*. have demonstrated that PPARδ orchestrates a pro-metastatic metabolic response to microenvironmental cues in pancreatic cancer. This metabolic rewiring promotes invasiveness and metastasis, and inhibition of PPARδ suppresses these effects^16^.

Our clinical observations of PPARδ expression correlating with adverse features and potentially poorer outcomes are consistent with these mechanistic insights. If these findings in a small series of 25 patients are confirmed, they will be important in reinforcing the role of PPARδ in driving aggressive disease biology through metabolic reprogramming and highlighting its potential as both a prognostic biomarker and a therapeutic target in PDAC.

## Conclusions

PPARδ immunohistochemistry in pancreatic cancer resections appears associated with adverse pathological features and poorer outcome. This small exploratory study highlights PPARδ as a potential biomarker of aggressive disease, meriting further investigation.

## Acknowledgements

The authors are grateful to the Comrie Cancer Research & Relief Club for financial support.

## Notes

### Competing Interest Statement

The authors have declared no competing interest.

